# Unsaturated Fatty Acids Are Required for Germline Proliferation and Membrane Structural Integrity in *Caenorhabditis elegans*

**DOI:** 10.1101/2025.07.24.666670

**Authors:** Bernabe Battista, Laura I. Lascarez-Lagunas, Diego de Mendoza, Monica P. Colaiácovo

**Affiliations:** Institute of Molecular and Cellular Biology of Rosario, National University of Rosario (IBR-CONICET-UNR) Ocampo y Esmeralda, Rosario 2000, Argentina; Department of Genetics, Blavatnik Institute, Harvard Medical School, Boston, MA 02115, USA

**Author notes:** **To whom correspondence should be addressed**: Diego de Mendoza, Monica P. Colaiácovo.

**Keywords:** *C. elegans*, inducible depletion of UFA, AID system, germline, fertility, meiosis, mitotic proliferation, meiotic progression, membrane structure

## Abstract

Unsaturated fatty acids (UFAs) are critical components of membrane lipids, but their specific roles in germline development and reproductive health remain poorly defined. Here, we investigated the consequences of UFA depletion in the *Caenorhabditis elegan*s germline using an auxin-inducible degron (AID) system to conditionally degrade FAT-7, the major Δ9 stearoyl-CoA desaturase, in a *fat-5; fat-6* double mutant background. This strategy bypassed the lethality associated with complete loss of Δ9 desaturase activity, enabling analysis of UFA deficiency in adult animals. UFA depletion led to a dramatic reduction in brood size, elevated embryonic and larval lethality, and a severe loss of germline nuclei. We found that UFAs are essential for mitotic proliferation, DNA replication, and chromosome organization in the germline. Moreover, reduced UFA levels impaired meiotic progression, accompanied by loss of membrane integrity in the syncytial germline. Notably, UFA deficiency increased nuclear pore complex (NPC) signal intensity, suggesting alterations to the nuclear envelope (NE). Together, our findings demonstrate that UFAs are indispensable for germline maintenance, affecting cell cycle progression, chromosome organization, and membrane architecture. These results underscore a fundamental link between acyl chain composition and reproductive success, highlighting the critical role of lipid homeostasis in the germline.

**Summary:** Unsaturated fatty acids (UFAs) are essential for fertility, but their role in germline maintenance remains unclear. In this study, we used *Caenorhabditis elegans* to examine how UFA depletion affects the germline. By conditionally disrupting UFA synthesis, we found that low UFA levels impair germline mitotic proliferation, DNA replication, meiotic progression, and germline membrane structures. These findings demonstrate that lipid composition is critical for germline maintenance and highlight a broader role for fatty acids in reproductive health, offering insights relevant to metabolic and fertility disorders in humans.

## Introduction

Globally, approximately 17.5% of the adult population, or one in six people, experience infertility (World Health Organization 2024). An increasing body of work suggests the existence of a positive correlation between lipid metabolism disorders—such as obesity, dyslipidemia, and metabolic syndrome—and reproductive problems (Robker 2008; Broughton & Moley 2017; Silvestris et al. 2018; Gambineri et al. 2019; Cohen et al. 2021; Yang et al. 2022). Moreover, lipid abnormalities affect approximately 45% of women of reproductive age (Pugh et al. 2017). Particularly, emerging evidence highlights the significant impact of fatty acid composition on female reproduction. The accumulation of saturated fatty acids (SFA), such as palmitic acid (PA 16:0) and stearic acid (SA 18:0), has been associated with several detrimental effects, including apoptosis in ovarian granulosa cells, restricted embryo development, impaired trophoblast migration, early placental development issues, and inhibited follicular growth and antrum formation (Mu et al. 2001; Jungheim et al. 2011; Wu et al. 2012; Valckx et al. 2014; Rampersaud et al. 2020). In contrast, the presence of unsaturated fatty acids (UFA) has been linked to positive reproductive outcomes. Oleic acid (OA 18:1) has been shown to support oocyte development and counteract the harmful effects of palmitic acid (Fayezi et al. 2018; Yousif et al. 2020; Zarezadeh et al. 2021). Similarly, arachidonic acid (AA 20:4) and eicosapentaenoic acid (EPA 20:5) play key roles in supporting oocyte maturation, embryonic development, and embryo implantation (Jawerbaum & Capobianco 2011; Vilella et al. 2013; Khajeh et al. 2017).

While there is a clear connection between fatty acid metabolism and reproductive health, the mechanisms through which fatty acids influence reproduction and germline maintenance remain poorly understood. In this study, we aim to investigate this interaction using *Caenorhabditis elegans*, a well-established model organism for studying lipid metabolism and sexual reproduction. *C. elegans* is a genetically and cytologically tractable model system that offers numerous advantages for such studies, including a well-defined and characterized germline, a rapid life cycle, and a high degree of shared gene and biochemical pathway conservation with humans (60%–80%) making it highly predictive of mammalian outcomes (Colaiácovo 2006; Lui & Colaiácovo 2013; Allard et al. 2013; Hillers et al. 2017). Therefore, *C. elegans* is a useful experimental system for uncovering the molecular mechanisms linking lipid metabolism and germline maintenance, offering insights for subsequent targeted mammalian studies.

In *C. elegans*, as in most eukaryotes, the synthesis of UFAs relies on the desaturation of SFA, a process initiated by Δ9 desaturases (Brock et al. 2006; Brock et al. 2007). This involves the genes *fat-6* and *fat-7*, which encode Δ9 stearoyl-CoA desaturases, and the gene *fat-5*, which encodes a Δ9 palmitoyl-CoA desaturase. The UFA synthesis pathway in *C. elegans* begins with PA, which is either derived from the *E. coli* diet or synthesized *de novo*. PA is converted to palmitoleic acid (16:1 Δ9) by the enzyme FAT-5. This product is subsequently elongated to cis-vaccenic acid (18:1 Δ11), an abundant fatty acid found in phospholipids and triglycerides. Alternatively, palmitic acid can be elongated to stearic acid (18:0), which serves as the substrate for FAT-6 and FAT-7, converting it into oleic acid (18:1 Δ9), an essential monounsaturated fatty acid (MUFA). OA is the substrate for the biosynthesis of all polyunsaturated fatty acids (PUFAs) the nematode possesses (Brock et al. 2007; Watts y Ristow 2017).

Here, we assess the impact of unsaturated fatty acid (UFA) deficiency on reproductive health and germline maintenance in *C. elegans*. To circumvent the lethality observed when all Δ9 desaturases are mutated in *C. elegans*, we employed the auxin-inducible degron (AID) system to generate a conditional *fat-7* gene, which encodes the major stearoyl-CoA desaturase (SCD), in a line where *fat-5* and *fat-6* are deleted (Battista et al. 2025). We show that UFA depletion results in a significantly reduced brood size, increased embryonic lethality, and a severe decrease in the number of germline nuclei. We find that UFA deficiency decreases mitotic proliferation and disrupts germline DNA replication, resulting in transient S-phase arrest. Moreover, UFAs are indispensable for proper meiotic progression, chromosome morphogenesis and maintaining membrane integrity of the syncitial structure in the *C. elegans* germline. Our study also uncovers a previously unrecognized role of UFAs in the establishment of nuclear pore complexes, emphasizing their fundamental contribution to nuclear organization. Collectively, these results underscore the need of maintaining a balanced SFA/UFA ratio in the germline and illuminate the broader significance of UFAs in reproductive health and cellular architecture.

## Materials and Methods

### Genetics and growth conditions

N2 Bristol worms were used as the wild-type background. Unless otherwise noted, lines were cultured under standard conditions as in Brenner (1974). The following mutations and chromosome rearrangements were used in this study: *ieSi57* (Zhang et al. 2015), *unc-119(ed3)* (Zhang et al. 2015), *fat-6(tm331)* (Brock et al. 2007)*, fat-7(dme1[fat-7::3xflag::aid])* (Battista et al. 2025), and *fat-5(tm420)* (Brock et al. 2007).

### Auxin-mediated protein degradation

Auxin-mediated degradation of FAT-7 in *fat-6*; *fat-5* double mutants was carried out by placing DDM6 worms on seeded NGM plates supplemented with 4mM auxin (dissolved in ethanol) from L3 to 24 hr post-L4 as described in Battista et al. 2025. Control plates were prepared by adding the same amount of ethanol and hexane as in the auxin-containing NGM plates. All plates were vented for about 1.5-2 hours for solvent evaporation, then stored at 4°C protected from light and used within a week. Ethanol exposure can induce dose-dependent physiological changes in *C. elegans*, including alterations in body length, maturation time, brood size, and lifespan (Davis et al. 2008; Alaimo et al. 2012), although *C. elegans* can develop ethanol tolerance over time (San-Marina et al. 1989; Kwon et al. 2004; Mathies et al. 2015). To ensure clear and reliable comparisons, we used ethanol-exposed worms as the primary control, while non-exposed worms served as additional controls for phenotype evaluation (Figure S1 a-f).

### Scoring embryonic lethality, larval lethality, frequency of males and brood size

To assess embryonic lethality, larval lethality, male frequency, and brood size, age-matched worms, grown in either auxin-containing or control NGM plates from the L3 to 24 hours post-L4 stage, were transferred to fresh NGM plates (containing either auxin, auxin+OA, or ethanol and hexane) every 24 hours over a period of four days. The total number of fertilized eggs laid, hatched, and the number of progeny (hermaphrodites and males) that reached adulthood were scored. At least two independent biological replicates were assessed for each condition.

### Immunofluorescence and imaging methods

*C*. *elegans* gonads from 24 hour post-L4 hermaphrodites were dissected and whole-mounted on slides as in Colaiácovo et al. (2003) using 4% paraformaldehyde fixation. Primary antibodies were used at the following dilutions: rabbit α-phospho-histone H3 (Ser10) (1:2000; Cell Signaling); rabbit α-SYX-4 (1:300, (Jantsch-Plunger y Glotzer 1999)); guinea pig α-pSer8 SUN-1 (1:700; (Woglar et al. 2013); mouse α-NPC (1:500) (Mab414 (ab24609), Abcam); and mouse α-BrdU (1:500) (BD). Secondary antibodies were purchased from Jackson ImmunoResearch Laboratories (West Grove, PA) as AffiniPure IgG (H+L) with minimum crossreactivity: α-rabbit Cy3 (1:200); α-guinea pig Alexa 488 (1:500); and α-mouse Alexa 488 (1:500). High-resolution imaging was performed using a IX-70 microscope (Olympus, MA) with a cooled CCD camera (CH350; Roper Scientific, AZ) driven by the DeltaVision Imaging System (Applied Precision, GE Healthcare). Images were collected at 0.2μm intervals using a 100x objective (N.A. 1.4), encompassing seven 36 x 36 μm zones starting from the distal tip, then deconvolved using a conservative ratio and 15x cycles with SoftWorx 3.3.6 software (Applied Precision), and processed with Fiji ImageJ (Schindelin et al. 2012).

General gonad morphology and evaluation of oocytes at diakinesis was performed in 24 hour post-L4 whole worms fixed with Carnoy’s fixative (6 ethanol:3 chloroform:1 glacial acetic acid) as in Villeneuve (1994) with some minor modifications. Specifically, fixed worms were rehydrated in 20 μl of M9 for 1 hour, excess M9 was then wicked away with a Whatman paper strip and 10 μl of a staining solution was added containing 50% DAPI for DNA visualization and 50% Vectashield (Vector Laboratories; Burlingame, CA) anti-fading agent. Images were captured using a ZEISS LSM880 laser scanning confocal microscope, employing a 60X objective with an additional 1.5X magnification.

### S-phase assessment by BrdU staining

S-phase was assessed using 5-bromo-2’-deoxyuridin staining (BrdU) in a 1:2.5 (BD) dilution, as described in Crittenden et al. (2006). Gonad dissections, immunostaining, antibody dilutions and imaging conditions were as described above. At least 10 gonads were scored for each condition at each timepoint analyzed from two independent biological repeats.

### Germ cell apoptosis

Germ cell corpses were scored using acridine orange staining, as described in (Kelly et al. 2000). A ZEISS LSM880 laser scanning confocal microscope was employed to visualize the apoptotic corpses along the gonad arm. The germlines of more than 90 worms from at least three independent biological repeats were scored for each condition/genotype (see Figures S2a, S2b and Table S1).

### Statistical methods

Statistical analyses were performed using GraphPad Prism version 8.0. Details of the statistical tests applied are provided in the corresponding figure legends. Briefly, the Fisher exact test was used to assess statistical significance for embryonic lethality, larval lethality and the frequency of male progeny (Him phenotype). The two-tailed unpaired *t*-test, 95% C.I. was applied to brood size data since here proportion comparisons between categories cannot be made. Group comparisons for germ cell apoptosis were carried out using the two-tailed Mann-Whitney test, 95% C.I.

## Results

### Reduction in UFA levels results in decreased brood size, increased embryonic and larval lethality, and germline defects

UFAs are essential for development in most eukaryotes, including *C. elegans*. Complete depletion of these fatty acids arrests growth, preventing the animals from reaching adulthood (Book et al., 2006; Brook et al., 2007). To bypass this limitation and assess the role of UFAs in the adult germline, we utilized a line herein referred to as DDM6: a loss-of-function *fat-6*; *fat-5* double mutant carrying a CRISPR-Cas9 engineered C-terminal AID-tagged *fat-7* for auxin-mediated degradation of this remaining Δ9-desaturase (Battista et al. 2025). Synchronized L1 larval stage DDM6 animals were placed on regular NGM plates until they reached the L3 stage (36h at 20°C). Animals were then moved to plates containing either vehicle alone (+EtOH), auxin (+AUX), or auxin + oleic acid (+AUX+OA) until they reached adulthood (24h-post-L4, ∼ 48h).

To assess the role of UFAs in fertility, we analyzed the total number of eggs laid (brood size), number of unhatched eggs (embryonic lethality), larval lethality, and the incidence of male progeny following UFA depletion. We observed a 2.5-fold reduction in brood size, indicative of sterility, upon UFA depletion compared to vehicle alone (Figures 1a, S1a and Table S1). This was accompanied by a 7-fold increase in embryonic lethality, with 30.2% embryonic lethality in auxin-treated worms compared to 4.4% in control animals (p < 0.0001 by the Fisher’s exact test; Figures 1a and S1a), and a 30-fold increase in larval lethality (p < 0.0001), with 93.6% in auxin-treated worms compared to 3.1% in controls, but no significant changes in the frequency of male progeny (p = 0.4162). As expected from the anticipated biosynthesis block, the reduced brood size and increased embryonic and larval lethality induced by auxin were rescued upon OA supplementation (p < 0.001, p < 0.0001, p < 0.0001; Figures 1a and S1a).

**Figure 1.**
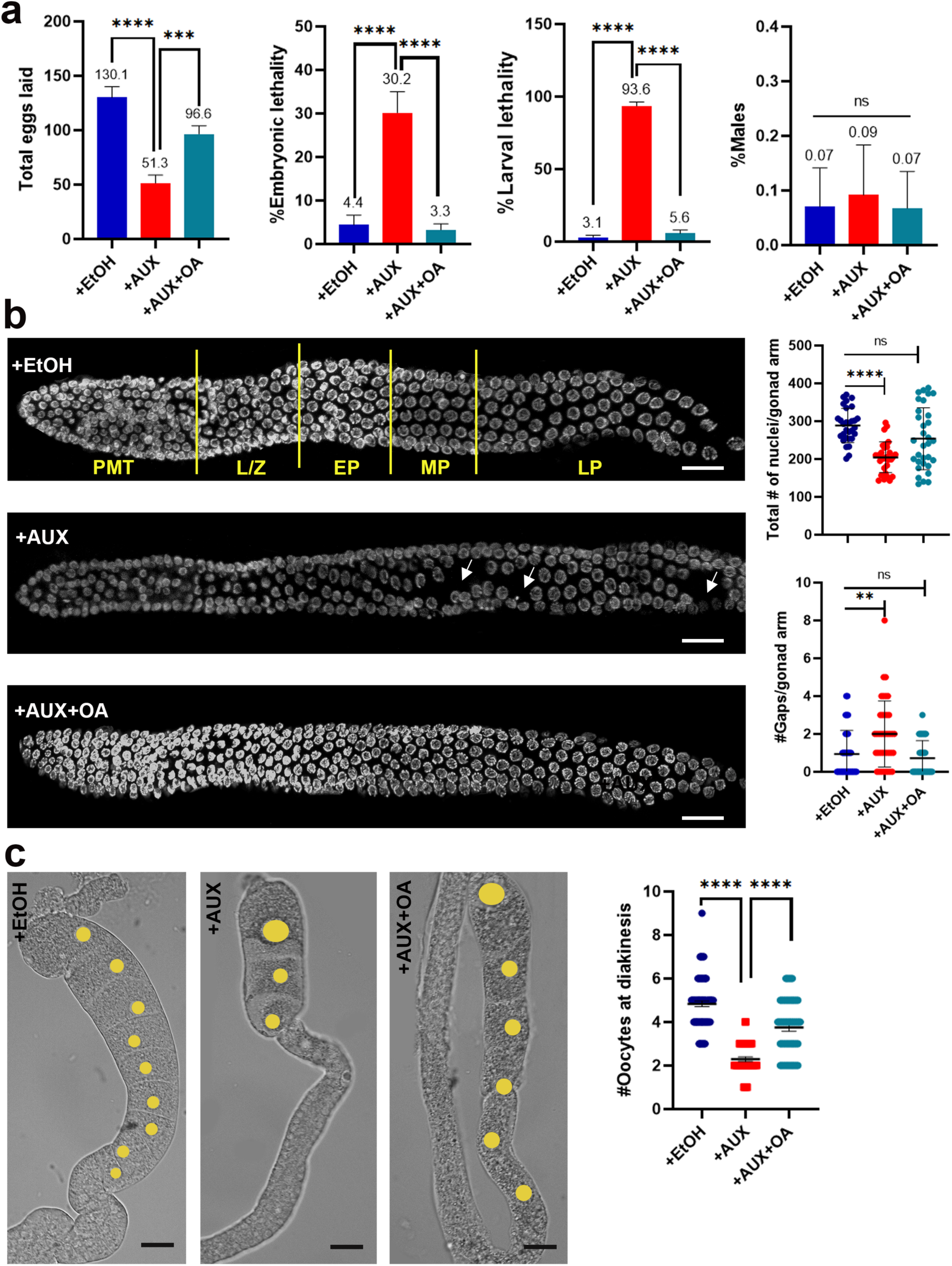
Reduction in UFA levels result in decreased brood size, increased embryonic and larval lethality, and germline defects. **(a)** Mean number of eggs laid (brood size), the percentage of embryonic lethality, larval lethality, and male progeny are shown for DDM6 worms grown on plates with ethanol (+EtOH), auxin (+AUX), or auxin plus oleic acid (AUX+OA) at 20°C. Error bars represent SEM. In the case of brood size, p values were calculated by the unpaired *t*-test, 95% C.I. ***p < 0.001, ****p < 0.0001. A minimum of 11 P0 worms were analyzed for each condition from two biological repeats. For embryonic lethality, larval lethality and male progeny: ****p < 0.0001, ns: non-significant, by the Fisher’s exact test, 95% C.I. **(b)** Depletion/reduction in UFA levels induces a significant reduction in the gonad size and increases the number of gaps (discontinuities or spaces lacking germ cell nuclei) per gonad arm. Left, images show DAPI stained gonads from Carnoy’s fixed whole worms exposed to the indicated conditions. Arrows indicate gaps in the gonad. PMT: pre-meiotic tip, L/Z: leptotene/zygotene, EP: early pachytene, MP: middle pachytene and LP: late pachytene. Scale bars, 20μm. Right, graph shows quantification of the total number of nuclei (top) and gaps (bottom) per gonad arm. Error bars represent SD. **p < 0.01, ****p < 0.0001, ns: non-significant by the two-tailed Mann-Whitney test, 95% C.I. A minimum of 30 gonads from at least 4 biological repeats were scored for each condition. **(c)** UFA depletion results in a decreased number of oocytes at diakinesis. (Left) Representative images of dissected gonads from DDM6 worms exposed to the indicated conditions. The yellow circles highlight each oocyte nucleus. Scale bars, 25μm. (Right) Graph shows quantification of the total number of oocytes at diakinesis. Error bars represent SEM. ****p < 0.0001 by the two-tailed Mann-Whitney test, 95% C.I. A minimum of 48 gonads from at least 3 biological repeats were scored for each condition.

Given the significant reduction in brood size observed for the UFA depleted worms, we investigated whether perturbations to germline nuclei contributed to the observed infertility. Germline nuclei are arranged in a spatial and temporal gradient, starting with nuclei undergoing mitotic proliferation at the distal tip region (referred to as the proliferative zone or pre-meiotic tip -PMT-), which then enter and progress through meiosis (Figure 1b; Kimble & Crittenden 2005). Analysis of DAPI-stained whole-mounted gonads revealed a significant reduction in the number of germline nuclei in auxin-treated gonads compared to vehicle alone (288 ± 45 and 204 ÷ 40, respectively; p < 0.0001, Mann-Whitney U-test, 95% C.I.; Figures 1b and S1b). This was accompanied by alterations in chromosome organization as indicated by a 2-fold increase in regions with reduced nuclear density (gaps) in auxin-treated compared to control gonads (0.93 ± 1.24 and 2 ± 1.74, respectively; p < 0.01; Mann-Whitney U-test, 95% C.I., Figures 1b and S1b). Both the decrease in total number of germline nuclei and the increase in gaps were fully rescued following OA supplementation (p = 0.055; p = 0.6301). In addition, UFA depletion also resulted in a reduction in the number of diakinesis-stage oocytes compared to controls (4.79 ± 1.06 and 2.29 ± 0.71, respectively; p < 0.0001, Mann-Whitney U-test, 95% C.I. Figure 1c). Although the auxin + OA condition also showed fewer diakinesis-stage oocytes compared to controls, the number was significantly higher than in the auxin-only condition (p < 0.0001, Mann-Whitney U-test, 95% C.I.; Figure 1c), indicating that fatty acid supplementation partially suppresses this auxin-induced phenotype. Overall, these findings suggest that reduced UFA levels result in decreased brood size, increased embryonic and larval lethality, a decrease in the number of germline nuclei, and defects in germline chromosome organization.

### Reduced UFA synthesis induces mitotic alterations

A reduction in the number of germline nuclei may have different origins including decreased mitotic proliferation, elevated germline apoptosis rates, and/or an accelerated oocyte maturation rate. Given the observed reduction in egg production, and the decreased number of diakinesis oocytes observed following auxin exposure, an increased oocyte maturation rate seems unlikely. To assess whether germline apoptosis contributes to the reduced number of germline nuclei, we evaluated the rates of programmed cell death in the germline by acridine orange staining as in Kelly et al. (2000). In *C. elegans*, physiological germ cell death is typically marked by the presence of 0 to 3 germ cell corpses near the gonadal bend during the late pachytene stage of meiosis (Gumienny et al. 1999; Gartner et al. 2000). Our observations revealed that auxin exposure did not increase apoptosis rates. Across all conditions tested, a mean between 0.04 to 0.4 germ cell corpses were detected in the conditions tested (Figures S2a, S2b and Table S1), yet no statistical differences were found, ruling out apoptosis as the cause of the reduced number of germline nuclei.

We next evaluated if mitotic proliferation might be altered in the gonad upon UFA depletion by scoring the number of nuclei in the PMT that were positive for histone H3 phosphorylation, a marker indicative of M phase (Seidel & Kimble 2015). Young adult animals (24 hours post L4) were dissected and their whole mounted gonads were immunostained for phospho-histone H3 (pH3). We found that the number of pH3-positive nuclei was significantly decreased in auxin-exposed animals compared to those in the ethanol control (1.0 and 2.75, respectively; p < 0.001 by the Mann-Whitney U-test, C.I. 95%; Figures 2a and S1c). Although the auxin +OA condition did not show a significant increase in pH3-positive nuclei compared to auxin-exposed worms (p = 0.2177), a trend toward a potential rescue was observed. The decreased number of pH3-positive nuclei suggests that UFAs are required for normal mitotic proliferation.

**Figure 2.**
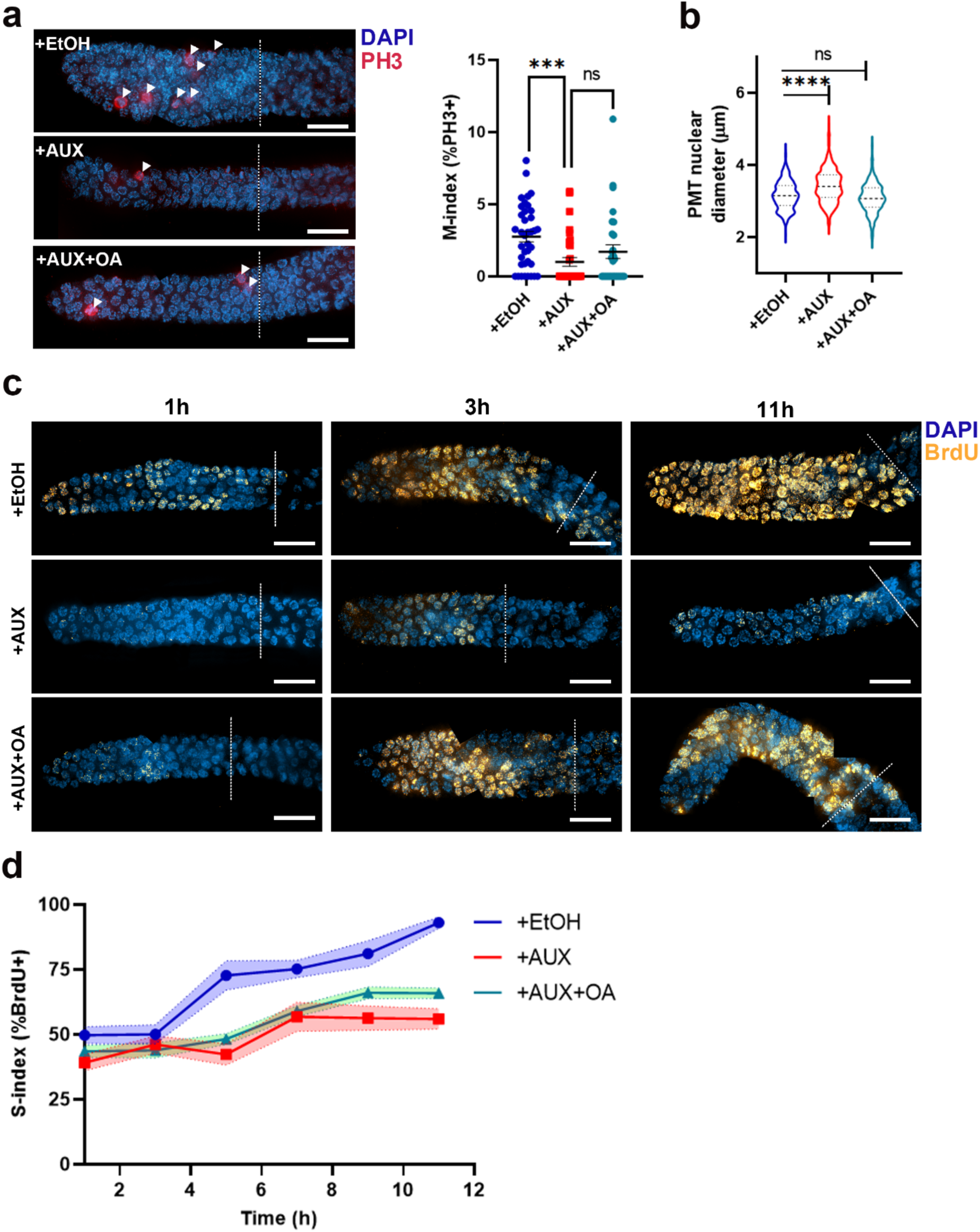
Reduced UFA synthesis induces mitotic alterations. **(a)** Auxin exposure causes altered mitotic proliferation in the *C. elegans* germline. Left, representative high-resolution images of nuclei in the pre-meiotic tip region (PMT) of the germline immunostained for phospho-histone H3 (Ser10) (pH3). The dashed line represents the PMT - L/Z (leptotene/zygotene) boundary and the arrows indicate pH3-positive nuclei. Scale bars, 15 μm. Right, graphs show quantification of M-index as percentage of pH3-positive nuclei in the PMT. Error bars represent SEM. ***p < 0.001, ns: non-significant by the two-tailed Mann-Whitney test, 95% C.I. A minimum of 30 gonads from at least 2 biological repeats were scored for each condition. **(b)** Distribution of the average nuclear diameter in the PMT represented by a violin plot. The bold dashed line indicates the median, while the upper and lower dashed lines correspond to the 75th and 25th percentiles, respectively. ****p < 0.0001 and ns: non-significant by the two-tailed Mann-Whitney test, 95% C.I. Between 180 and 400 individual nuclei diameters were scored for each condition from 3 biological repeats. **(c)** Reduction in UFA levels slows down S-phase. Representative high-resolution images from mounted gonads of worms exposed to the indicated conditions and co-stained with DAPI after BrdU pulses of increasing times. The dashed line represents the PMT - L/Z boundary. Scale bars, 15 μm. **(d)** Time course for complete incorporation of BrdU in the PMT region. The graph depicts the percentage of BrdU-positive nuclei within the PMT at different times of incubation for the different conditions. The faded-colored area depicts error bars as SEM.

Further analysis revealed an increase in nuclear diameter at the PMT following UFA depletion relative to control (3.42 µm and 3.16 µm, respectively; p < 0.0001, Mann-Whitney U-test, C.I. 95%; Figures 2b and S1d). An enlargement of the nuclei in the proliferative zone can stem from activation of a replication-dependent S-phase checkpoint resulting in transient S-phase arrest (Chen et al. 2003; Bartek et al. 2004; Uto et al. 2004; Garcia-Muse y Boulton 2005).Therefore, we assessed the S-phase index (percentage of nuclei in S-phase) by labeling with BrdU (5-bromo-2’-deoxyuridine), a thymidine analog which is incorporated into cells undergoing active DNA replication (Crittenden et al. 2006). Young adult hermaphrodites (24 h post L4) were exposed to BrdU-labeled *E. coli* MG1693 for 1, 2, 3, 5, 7, 9, and 11 hours (Figure 2B, C and Supplemental Table 1). 49%, 50% and 93% of PMT nuclei were labeled after 1, 3 and 11 hours, respectively, in control (+EtOH). In contrast, the S-index in the PMT was significantly decreased at 1 and 11 hours (39% and 56%, respectively; p < 0.05 and p < 0.0001), but not at 3 hours (46%; p = 0.437 by the unpaired *t*-test, 95% C.I.), following auxin-mediated depletion (Figures 2c, 2d and S1e). Supplementation with OA partially restored the S-index at 1 and 11 hours (43% and 66%, respectively). These results indicate that reduced UFA synthesis can interfere with DNA replication, potentially slowing this process in the germline. Taken together, these data suggest that the reduction in the number of nuclei in the UFA depleted gonads results in part from decreased mitotic proliferation and S-phase checkpoint activation resulting in a reduction in DNA replication rate.

### UFAs are required for normal meiotic progression

Analysis of DAPI-stained whole-mounted gonads revealed that 86.7% of gonads from auxin-exposed worms had nuclei at the pachytene stage with chromosomes arranged in a crescent-shaped configuration, characteristic of earlier leptotene/zygotene stage, compared to 20% and 13.3% in control and OA supplementation conditions, respectively (p < 0.001 and p < 0.0001) by the two-sided Fisher’s exact test; n=15 gonads each) (Figure 3a). This pattern is indicative of the presence of lagging leptotene/zygotene nuclei within the later stages of meiotic prophase I, which could be suggestive of altered meiotic progression (Cuenca et al. 2020). To evaluate potential disruptions in meiotic progression upon UFA depletion, we next analyzed the distribution of phosphorylated SUN-1 (S8 pSUN-1) (Figure 3b). Under normal conditions, pSUN-1 exhibits intense clustering at the nuclear envelope (NE) during leptotene/zygotene and transitions to a weaker, more dispersed signal within the nucleus as cells progress to early and mid-pachytene (Woglar et al. 2013). To account for the size differences observed between gonads of auxin-exposed worms and control worms, we quantified the number of rows in which at least two nuclei were positive for pSUN-1 within a row and normalized this to the total number of nuclei in the gonad arm, thereby excluding size-based variability. We found a significant increase in the number of normalized pSUN-1-positive rows in auxin-exposed worms compared to the control group (13.84 vs. 8.45, respectively; p < 0.0001, unpaired *t*-test, 95% C.I., Figures 3b and S1f), which was partially rescued by OA supplementation (10.60; p < 0.001). These observations show that reduced UFA synthesis induces meiotic progression defects in the germline. However, the lack of increased germ cell apoptosis in late pachytene (Figure S2 A, B) and the presence of six intact DAPI-stained bodies in the -1 and -2 oocytes at diakinesis, representing the six pairs of attached homologs (bivalents) across all the conditions assayed (Figures S2c and S2d), suggest that the defect in meiotic progression upon reduction in UFA content does not stem from problems with either chromosome synapsis or DNA double-strand break repair during meiosis, which would have led to increased apoptosis and univalents at diakinesis.

**Figure 3.**
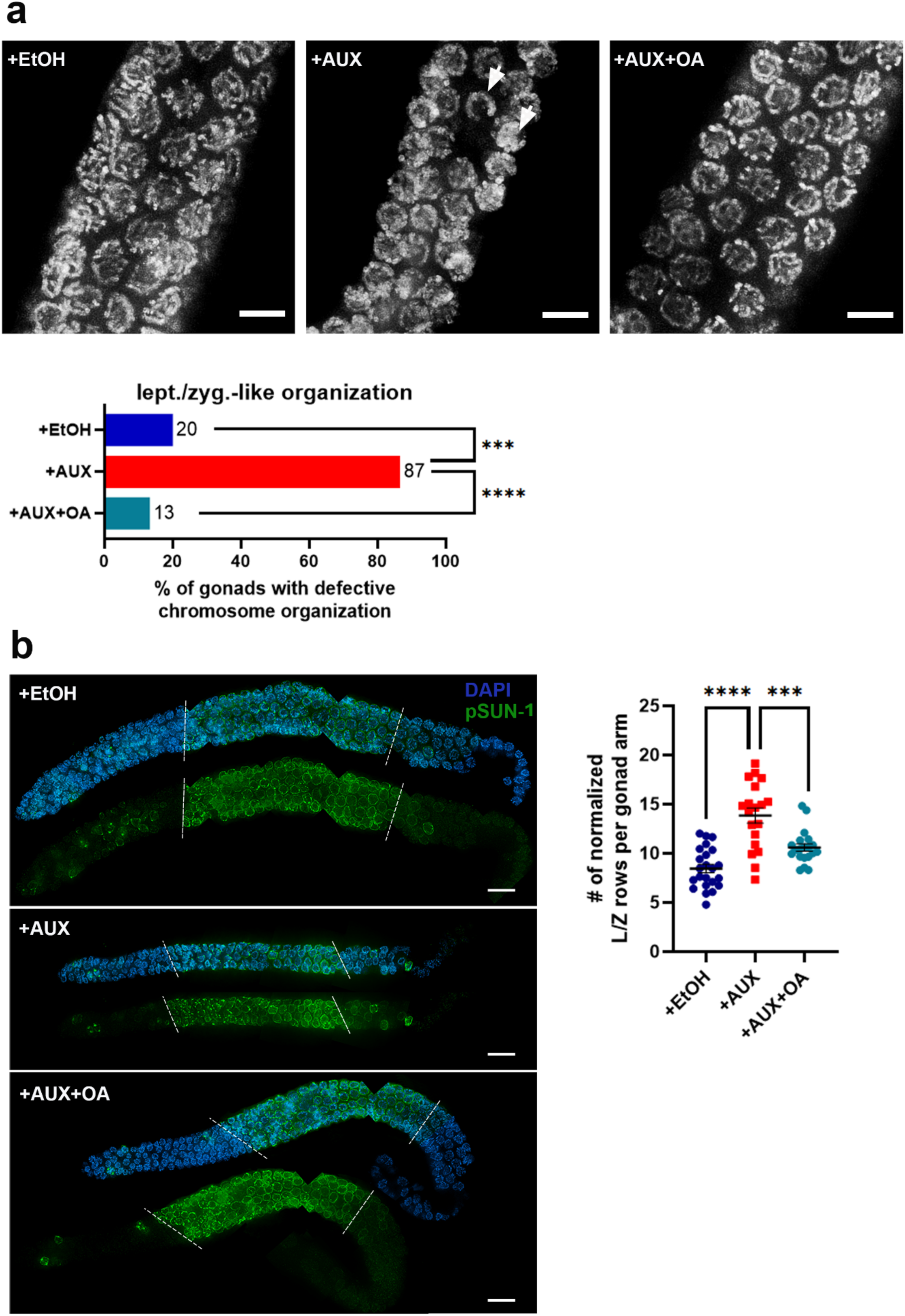
UFAs are required for normal meiotic progression. **(a)** Top, Images of pachytene nuclei from dissected gonads fixed with 4% paraformaldehyde and stained with DAPI. A minimum of 13 gonads were scored for each indicated condition from at least 2 biological repeats. Arrows indicate nuclei with chromatin in leptotene/zygotene-like organization. Scale bars, 5 μm. Bottom, graph shows the percentage of gonads with nuclei exhibiting chromatin in a leptotene/zygotene-like organization at the pachytene stage. ***p < 0.001, ****p < 0.0001 by the two-sided Fisher’s exact test (bottom). **(b)** Left, high-resolution images of whole mounted gonads of DDM6 worms exposed to the indicated conditions co-stained with anti-S8 pSUN-1 (green) and DAPI (blue). Dashed lines demarcate the first (left) and the last (right) complete row with nuclei showing S8 pSUN-1 signal. Scale bars, 15 μm. Right, graph shows the number of normalized rows in leptotene/zygotene (L/Z) stage (pSUN-1-positive) per gonad for each condition. Error bars represent SEM. ***p < 0.001, ****p < 0.0001 by the two-tailed unpaired *t*-test, 95% CI. Between 19 and 23 gonads for each condition from two independent biological repeats were analyzed. The detection of some pSUN-1-positive nuclei in the premeiotic tip is consistent with previous reports of pSUN-1 nuclei being observed in the premeiotic tip of wild-type worms, as well as in normal mitotic nuclei in *C. elegans* embryos (Woglar et al. 2013; Zuela & Gruenbaum 2016; Shin et al. 2019).

### Alterations in lipid homeostasis disrupt germline membrane structures in a manner that is reversible by oleic acid supplementation

A plausible explanation for the defects observed in the germline due to reduced UFA levels is the disruption of membrane structures. Phospholipids, the primary components of membranes, rely on their fatty acid composition to maintain the biophysical properties essential for proper membrane function (Watts y Ristow 2017; de Mendoza & Pilon 2019). Studies in *paqr-1; paqr-2* double mutants, known to accumulate high levels of saturated fatty acids in *C. elegans*, showed that low UFA levels can compromise the incomplete membranes surrounding germline nuclei before cellularization is completed at diakinesis (Devkota et al. 2021).

To assess the integrity of the germline syncytial structure upon UFA depletion, we analyzed the distribution of SYX-4, which localizes to the incomplete membranes separating germ cell nuclei (Jantsch-Plunger & Glotzer 1999). SYX-4 localizes in a honeycomb-like pattern tracking with the cell membrane structures within the germline as observed in wild type (N2) and untreated DDM6 (Figure 4a). While the hexagonal pattern was maintained in control (DDM6 + EtOH) worms, it was severely disrupted in auxin-exposed worms, with regions where the signal was completely lost. The signal was particularly altered in the mid and late pachytene regions (Figure 4a). Notably, supplementation with OA significantly suppressed the disruption caused by reduced UFA levels. Additionally, we quantified the intensity of the SYX-4 signal across the different regions of the gonad (Figure 4b and Table S2). In the DDM6 + EtOH control and the worms supplemented with OA, SYX-4 signal increased progressively from the premeiotic tip (PMT) to the late pachytene (LP) region. However, in auxin-exposed worms, the SYX-4 signal remained constant regardless of the region analyzed. This could be explained by SYX-4 localization being dependent on the varying lipid composition across different gonadal regions. In auxin-exposed worms, these region-specific lipid differences may be disrupted, preventing variations in SYX-4 signal localization as reflected by the lack of changes in signal intensity. However, it is important to consider the potential for nonspecific effects of ethanol exposure on the fluorescence signal. Comparisons across all conditions containing ethanol revealed higher fluorescence values, whereas the untreated DDM6 and N2 worms exhibited comparable fluorescence levels throughout the gonad.

**Figure 4.**
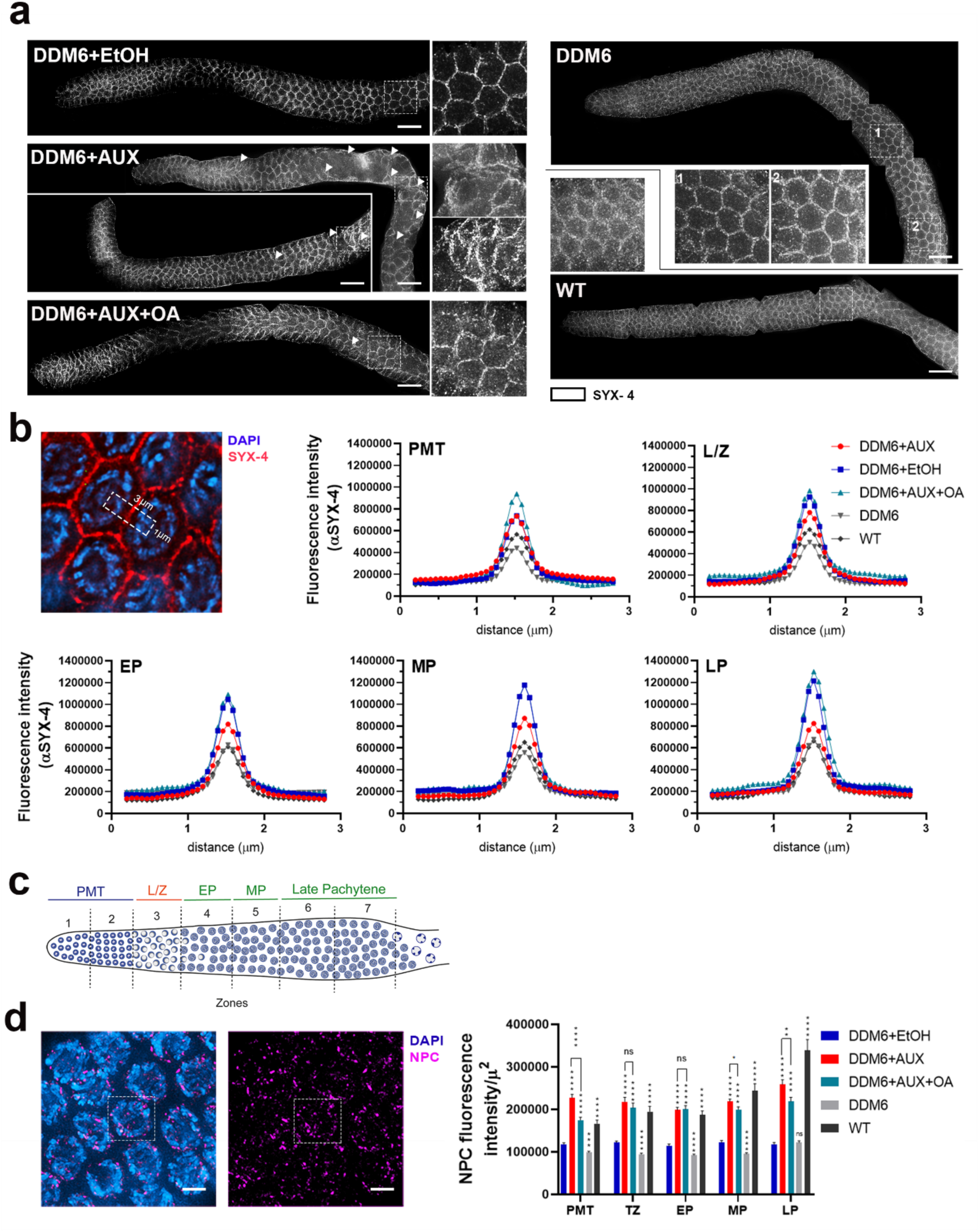
Low levels of UFAs alter cell membranes within the germline. **(a)** High-resolution representative images of whole mounted gonads of DDM6 worms exposed to the indicated conditions stained with anti-SYX-4. Images show the cell membrane honeycomb-like pattern observed for this marker which is perturbed to varying degrees upon UFA depletion. Dashed boxes indicate the selected areas shown at higher magnification. Arrowheads indicate regions where the honeycomb-like pattern is altered. Scale bars, 15 μm. **(b)** Left, high-resolution images of late pachytene nuclei (zone 6) from whole-mounted gonads stained with SYX-4 (red) and co-stained with DAPI (blue). An image capturing the mid-section of nuclei is shown. Right, quantification of SYX-4 signal intensity on each region of the gonad. Fluorescence intensity was measured from images captured through the center of the nuclei in a rectangular area (1 μm x 3 μm) that encompasses the border between two nuclei using Image J. Data represent average signal measured from at least 40 nuclei from five to six different gonads for each condition. **(c)** Top left, diagram of the *C. elegans* germline oriented from left to right. The different stages of meiotic prophase are indicated above the gonad and the seven equally sized zones evaluated are delimited by vertical dashed lines. Nuclei in zones 1 and 2 are undergoing mitosis (PMT) before entering meiosis at zone 3, corresponding to the leptotene/zygotene stages (L/Z). In zones 4-7 nuclei proceed through pachytene until they reach diplotene (EP, early pachytene; MP, mid-pachytene; LP, late pachytene). Bottom left, high-resolution images of late pachytene nuclei (zone 6) from whole-mounted gonads stained with NPC (magenta) and co-stained with DAPI (blue). Maximum intensity projections of all the z-stacks covering each nucleus are shown. Dotted square indicates the region where the intensity was measured. Intensity was adjusted to the average nucleus size of each region of the gonad. Scale bars, 3 μm. Bottom right, graph shows the mean number of NPC fluorescence intensity per gonad region for each condition. Data represents average normalized signal measured from 30 nuclei from six different gonads for each condition. *p < 0.05, **p < 0.01, ***p < 0.001, ****p < 0.0001 and ns: non-significant by the Mann-Whitney test, 95% C.I.

To test whether the nuclear envelope (NE) was affected under low UFA levels, we examined the distribution of nuclear pore complex (NPC) proteins (Figure 4c). While no differences in the spatial distribution of the NPC signal were observed across the conditions tested, we did detect a significant increase in signal intensity in the auxin-exposed condition compared to the control across all regions of the germline (p < 0.0001, Mann-Whitney U-test, C.I. 95%, Figure 4c). Notably, these changes were partially rescued by OA supplementation in the premeiotic tip, mid-pachytene, and late pachytene regions. We also observed that NPC fluorescence intensity is higher in *C. elegans* wild type (N2) worms compared to DDM6 worms in the absence of auxin treatment (Figure 4c). Notably, the DDM6 strain expresses only FAT-7, the major Δ9 desaturase in *C. elegans*, but lacks FAT-5 and FAT-6 desaturase activity (Battista et al. 2025). Although previous studies have suggested that the loss of either FAT-6 or FAT-7 can be compensated by the remaining desaturase, our findings indicate that subtle alterations in lipid homeostasis may influence NPC distribution. Collectively, these findings suggest that reduced UFA levels impair the formation or maintenance of membrane structures in the germline at both cellular and nuclear levels, as indicated by altered SYX-4 and NPC signals.

## Discussion

In the present study we have examined the role of UFAs in the *C. elegans* germline. Our results showed that a reduction in UFA levels significantly impacts brood size, embryonic and larval lethality but not the incidence of male progeny (Figures 1a and S1a). These findings align with previous studies indicating that UFAs are essential for proper development in *C. elegans*, as complete elimination of these compounds halts growth and induces sterility (Brock et al. 2006; Brock et al. 2007; Svensk et al. 2013; Devkota et al. 2021). Additionally, we found that the reduction in brood size is at least partially attributable to a reduction in the number of germline nuclei (Figures 1b and S1b). Further analysis revealed that this reduction is driven by diminished mitotic proliferation, rather than increased apoptosis or accelerated oocyte maturation, as evidenced by a significant decrease in phospho-histone H3-positive nuclei (Figures 2a and S1c). These observations are consistent with reports showing that Δ9 desaturation is needed to sustain ovarian cancer proliferation and tumor growth in xenomorphic models, as well as in glioblastoma stem cells (Li et al. 2017; Pinkham et al. 2019; Zhao et al. 2022).

Several studies have shown that cells require nutrients and lipids for successful cell division (Wymann & Schneiter 2008; Li et al. 2017; Zhao et al. 2022); however, the specific role of lipids in cell cycle regulation remains largely unexplored. In this study, we report a significant reduction in DNA synthesis rates in auxin-exposed worms compared to controls, as evidenced by decreased BrdU labeling (Figures 2c, 2d and S1e). Our findings suggest that fatty acid synthesis is rate-limiting for germline nuclei within the syncytial gonad to enter S phase when fatty acid availability is low. Consistent with this, a study using cell culture has identified lipid checkpoints during the G1 phase that prevent nuclei from progressing into S phase when lipid synthesis, including the production of UFAs, is insufficient (Köberlin et al. 2024). In the same study, it is noted that a shift from saturated to unsaturated lipids is essential for the G1/S transition and the continuation of cell cycle progression, as unsaturated lipid species remain elevated throughout the subsequent G2 and M phases. A plausible interpretation of our findings is that the depletion of unsaturated lipids (auxin exposure) disrupts this required lipid shift, thereby preventing germline nuclei from successfully completing mitosis. It is possible that mitogens must stimulate lipid accumulation and remodeling to enable germ cells to progress through the cell cycle. This mechanism would ensure an adequate supply of UFA components, supporting both membrane remodeling and DNA replication. Additionally, we observed an increase in nuclear diameter (Figures 2b and S1d), which may be linked to the activation of a replication-dependent S-phase checkpoint resulting in transient S-phase arrest (Boxem et al. 1999; Chen et al. 2003; Uto et al. 2004; van den Heuvel 2005). Together, these observations suggest that reduced UFA levels can disrupt cell cycle progression in the germline.

Meiotic progression was also disrupted, as evidenced by an increased number of rows of nuclei exhibiting phosphorylated SUN-1-positive signal in auxin-exposed worms compared to control (Figures 3 and S1f). A recent study in mouse testes has highlighted the importance of fatty acid desaturation in meiosis, where low UFA levels led to impaired telomere attachment to the nuclear envelope due to decreased membrane fluidity (Zhang et al., 2024). Since the protein machinery responsible for telomere-led chromosome movements during meiosis relies on the integrity of the nuclear envelope (NE) for proper attachment of the telomeres to the NE (Zetka et al. 2020), this finding underscores the significance of lipid composition in meiotic progression. Given the high conservation of these processes across species, it is likely that similar disruptions in chromosome dynamics occur in the *C. elegans* germline under conditions of reduced UFA availability.

The membrane disruptions observed in the germline of auxin-exposed worms may provide insights into how reduced levels of unsaturated fatty acids (UFAs) compromise fertility. The altered distribution of SYX-4 and increased NPC fluorescence intensity suggest that UFAs are essential not only for maintaining general membrane integrity and function but also for preserving the structural integrity of the NE. Supporting the idea that decreased lipid desaturation underlies these defects, the external supplementation of oleic acid, which restores polyunsaturated fatty acid (PUFA) synthesis, partially rescued the membrane and NE phenotypes. These findings are consistent with recent studies in yeast demonstrating that NE geometry, as well as NPC architecture, distribution, and function, are regulated by lipid unsaturation homeostasis (Romanauska y Köhler 2023). In this context, unsaturated lipids play a key role in reducing membrane rigidity and increasing fluidity due to their kinked acyl chains, which prevent tight packing of lipid tails. Maintaining a balanced composition of saturated and unsaturated lipids is crucial for proper pore membrane curvature and NPC integrity. Excessive lipid saturation can decrease membrane elasticity, leading to increased nuclear rigidity and a higher risk of fracturing.

Finally, it is worth noting that OA media supplementation was not always sufficient to fully or even partially suppress the deleterious phenotypes induced by auxin exposure. This is evident in the inability to restore the number of nuclei exhibiting pH3 signal, which indicates mitotic proliferation. These findings underscore the fundamental role of *de novo* UFA biosynthesis, demonstrating that exogenous dietary supplementation cannot always meet the internal demand for these compounds. Factors such as compound delivery to specific tissues or cellular locations, final intracellular concentration, compound metabolism, and overall availability may influence the phenotypic outcome (Gibson & Skett 1986). Moreover, it has been reported that newly synthesized fatty acids upon mitotic exit are used for phospholipid synthesis and incorporated mainly in the NE, suggesting they are required for its reassembly and/or expansion (Rodriguez Sawicki et al. 2019). Taken together, our findings highlight the essential role of UFAs in germline maintenance in *C. elegans*. Reduced UFA levels result in severe germline defects, disrupting mitotic and meiotic progression, and compromising membrane integrity, ultimately leading to decreased fertility. These insights deepen our understanding of the importance of lipid metabolism in reproductive biology with broader implications for germline maintenance in other organisms and human health.

## Data availability

Strains and plasmids are available upon request. The authors affirm that all data necessary for confirming the conclusions of the article are present within the article, figures, and tables. Table S1 contains detailed raw data for all the assessments performed except for SYX-4 determinations which are in Table S2. Supplemental material available online.

## Acknowledgements

WT N2 strain was kindly provided by the Caenorhabditis Genetics Center. We thank Dr. Verena Jantsch for the α-pSer8 SUN-1 antibody, Dr. Cecilia Vranych for assistance with *C. elegans* growth and maintenance, Viviana Villalba for excellent technical assistance, and Consejo Nacional de Investigaciones Científicas y Técnicas (CONICET) for the scholarship granted to

B.B. D.d.M is a member of the Carrera del Investigador Científico of CONICET.

## Funding

This research was supported by the National Institutes of Health grant R01GM072551 to M.P.C and by a Richard Lounsbery Foundation grant (2024-2025) to D.d.M.

## Conflicts of interest

The author(s) declare no conflicts of interest.

## Notes

### Competing Interest Statement

The authors have declared no competing interest.

