## Supplemental Material for "Unsaturated Fatty Acids Are Required for Germline Proliferation and Membrane Structural Integrity in *Caenorhabditis elegans*"

**Figure S1. Analysis of brood size, embryonic and larval lethality, and germline defects observed in untreated DDM6 (-EtOH) and wild type (N2) worms. (a)** Mean number of eggs laid (brood size), the percentage of embryonic lethality, larval lethality, and male progeny are shown for untreated DDM6 (-EtOH) and wild type (N2) worms. Error bars represent SEM. *p < 0.05 and ns, non-significant by the Fisher’s exact test, 95% CI. A minimum of 11 P0 worms were analyzed for each condition from two biological repeats. **(b)** Graph shows quantification of total number of nuclei (left) and gaps (right) per gonad. A minimum of 30 gonads from at least 4 biological repeats were scored for each genotype. Error bars represent SEM. ns: non-significant by the two-tailed Mann-Whitney test, 95% C.I**. (c)** Graph shows quantification of M-index as percentage of pH3-positive nuclei in the PMT. Error bars represent SEM. ns, non-significant by the two-tailed Mann-Whitney test, 95% C.I. A minimum of 30 gonads from at least 2 biological repeats were scored for each condition. **(d)** Mean nuclear diameter in the PMT. Error bars represent SEM. ns: non-significant by the two-tailed Mann-Whitney test, 95% C.I. 180 individual nuclei diameters were scored for each condition from 3 biological repeats. **(e)** Time course for complete incorporation of BrdU in the PMT region. Graph depicts the percentage of BrdU-positive nuclei within the PMT at different times of incubation for the indicated genotypes. The faded-colored area depicts error bars as SEM. **(f)** Graph shows the number of normalized rows in leptotene/zygotene (L/Z) stage per gonad (pSUN-1-positive) for each genotype. Error bars represent SEM. For DDM6, n=18, and for WT, n=15, where n represents the number of gonads scored. ns: non-significant by the two-tailed unpaired *t*-test, 95% C.I.

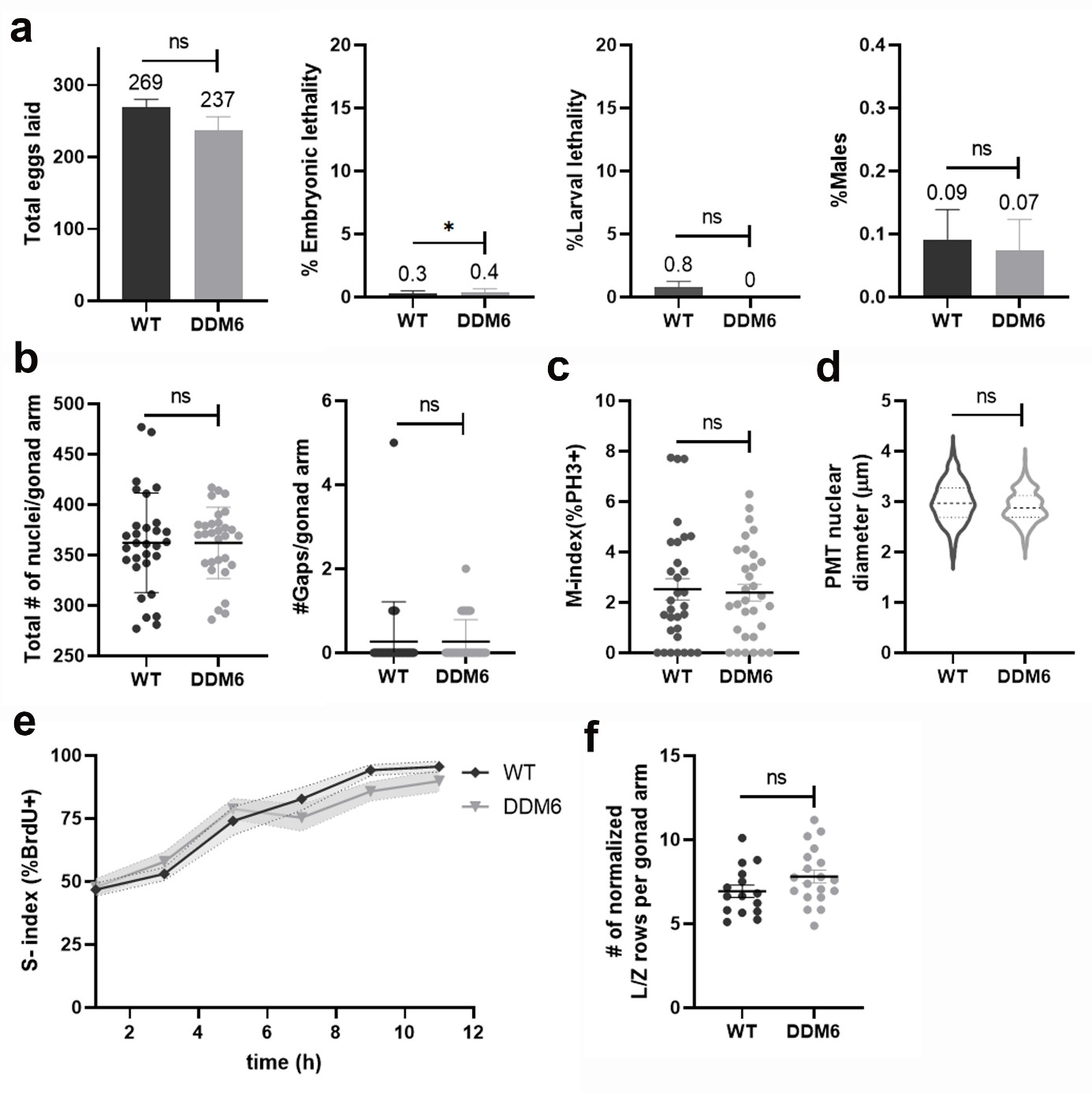

**Figure S2. Reduced UFA levels do not alter levels of germ cell apoptosis or the number of bivalents and chromosome morphology in late diakinesis oocytes. (a)** Representative images of acridine-orange-stained germ cell corpses (bottom) and the same germlines visualized by DIC optics (top). Arrows indicate germ cell corpses. Scale bars, 20 μm. **(b)** Graph shows mean number of germ cell corpses per gonad arm from three independent biological repeats, with more than 95 gonads scored for each condition. Error bars represent SEM. ns: non-significant by the two-tailed Mann-Whitney test, 95% C.I. **(c)** High-resolution representative images of DAPI-stained diakinesis nuclei. Images correspond to the oocyte positioned right before the spermatheca (-1 oocyte). Six intact bivalents are observed in oocytes from all assayed conditions. Each individual DAPI-stained body is indicated with a white number. Scale bars, 5 μm. **(d)** Quantification of the chromosome morphology defects observed in diakinesis oocytes. n = total number of oocytes scored. Lower case letters indicate the different categories evaluated and are described at the bottom of the table.

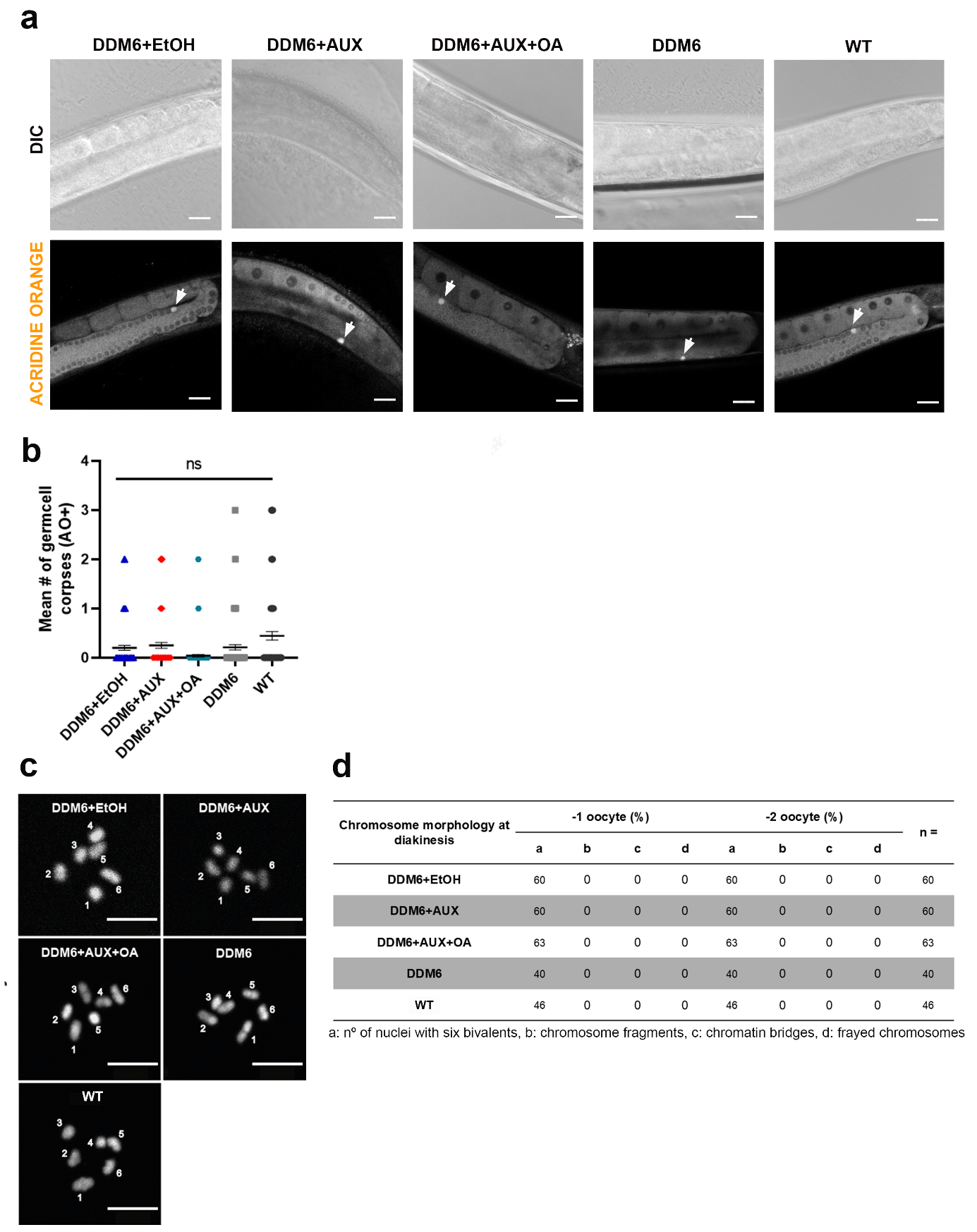
